# Atazanavir inhibits SARS-CoV-2 replication and pro-inflammatory cytokine production

**DOI:** 10.1101/2020.04.04.020925

**Authors:** Natalia Fintelman-Rodrigues, Carolina Q. Sacramento, Carlyle Ribeiro Lima, Franklin Souza da Silva, André C. Ferreira, Mayara Mattos, Caroline S. de Freitas, Vinicius Cardoso Soares, Suelen da Silva Gomes Dias, Jairo R. Temerozo, Milene Miranda, Aline R. Matos, Fernando A. Bozza, Nicolas Carels, Carlos Roberto Alves, Marilda M. Siqueira, Patrícia T. Bozza, Thiago Moreno L. Souza

**Author notes:** Correspondence footnote: Thiago Moreno L. Souza, PhD, Fundação Oswaldo Cruz (Fiocruz), Centro de Desenvolvimento Tecnológico em Saúde (CDTS), Instituto Oswaldo Cruz (IOC), Pavilhão Osório de Almeida, sala 16, Av. Brasil 4365, Manguinhos, Rio de Janeiro - RJ, Brasil, CEP 21060340. These authors contributed equally to this work (Natalia Fintelman-Rodrigues, Carolina Q. Sacramento, Carlyle Ribeiro Lima, Thiago Moreno L. Souza).

## Abstract

Severe acute respiratory syndrome coronavirus 2 (SARS-CoV-2) is already responsible for far more deaths than previous pathogenic coronaviruses (CoVs) from 2002 and 2012. The identification of clinically approved drugs to be repurposed to combat 2019 CoV disease (COVID-19) would allow the rapid implementation of potentially life-saving procedures. The major protease (Mpro) of SARS-CoV-2 is considered a promising target, based on previous results from related CoVs with lopinavir (LPV), an HIV protease inhibitor. However, limited evidence exists for other clinically approved antiretroviral protease inhibitors, such as atazanavir (ATV). ATV is of high interest because of its bioavailability within the respiratory tract. Our results show that ATV could dock in the active site of SARS-CoV-2 Mpro, with greater strength than LPV. ATV blocked Mpro activity. We confirmed that ATV inhibits SARS-CoV-2 replication, alone or in combination with ritonavir (RTV) in Vero cells, human pulmonary epithelial cell line and primary monocytes, impairing virus-induced enhancement of IL-6 and TNF-α levels. Together, our data strongly suggest that ATV and ATV/RTV should be considered among the candidate repurposed drugs undergoing clinical trials in the fight against COVID-19.

## 1) Introduction

Coronaviruses (CoVs) are single-stranded positive sense RNA viruses with large enveloped nucleocapsids that are able to infect a range of hosts including both animals and humans^1^. Although a number of human CoV are known to circulate seasonally, two highly pathogenic variants emerged in the 21^st^ century that cause life-threatening infection, the severe acute respiratory syndrome (SARS-CoV) and middle-east respiratory syndrome (MERS-CoV)^2^. At the end of 2019, a novel variant of SARS-CoV (SARS-CoV-2) appeared in the citizens of the City of Wuhan, China that is believed to have spilled over to humans from animal reservoirs, most likely bats and/or pangolins^3^. The novel 2019 CoV is phylogenetically closer to SARS-CoV (from the 2002 outbreak) than MERS-CoV (from 2012 outbreak) ^2,3^. Both SARS- and MERS-CoV raised international public health concerns with rates of mortality of 10 and 35%, respectively^4,5^. Soon after its discovery, the contemporary SARS-CoV-2 became a pandemic threat, with the number of confirmed infections ramping up globally^6^. To date, SARS-CoV-2 is responsible for 10 times more deaths than the total sum from SARS- and MERS-CoV, with more causalities daily that are continue to scale up^6^.

Currently, the most effective response to the SARS-CoV-2 pandemic has been self-quarantining and social distancing to avoid contact between infected and uninfected individuals that can flatten the virus dissemination curve, which aim to reduce the burden on medical resources to prevent loss of service for those with the highest need. While these social actions can disrupt virus transmission rates, they are not expected to reduce the absolute number of infected individuals. Furthermore, these strategies are also provoking a severe reduction in global economic activity^7^. To effectively combat the impact of SARS-CoV-2 on infected individuals, and society as a whole, it is essential to identify antiviral drugs for immediate use, as well as develop new drugs and a vaccine for long-term solutions to the disease associated with SARS-CoV-2 (COVID-19).

Repurposing of clinically approved drugs is the fastest pathway towards an effective response to a pandemic outbreak^8^. Some of the most promising antiviral candidates against SARS-CoV-2 have been under investigation since the outbreak of SARS-CoV in 2002. Building on this continuous investigation, an unprecedented effort to run a global clinical trial, called SOLIDARITY, is ongoing under the auspicious of the World Health Organization (WHO) and the United Nations (UN)^9^. This mega trial has been putting forward lopinavir (LPV)/ritonavir (RTV), in combination or not with interferon-β (IFN-β), chloroquine (CQ) and remdesivir to treat COVID-19^9^. LPV, RTV and remdesivir target viral enzymes, while the actions of CQ and IFN-β target host cells.

The most successful antiviral drugs often directly target viral enzymes^10^. For CoVs, its major protease (Mpro) has been a promissing drug target for almost two decades, starting with early studies on 2002 SARS-CoV that showed this enzyme to be inhibited by LPV/RTV, inhibitors of HIV protease^11^. Mpro is required during the CoV replication cycle to process viral polyprotein^12^. Highly pathogenic CoVs contain two open reading frames, ORF1a and ORF1b, that are translated by host ribosomes into their two respective viral polyproteins, pp1a and pp1ab. ORF1a encodes two cysteine proteases, the papain-like protease (PLpro) and Mpro. While PLpro cuts the polyprotein at three sites, Mpro is responsible for cleavage at 11 another locations that, together, produce the 16 nonstructural proteins.

In a combined therapy of LPV with RTV, LPV is included as the principle antiviral compound and RTV as an inhibitor cytochrome p450^13^. Although RTV can also display weak anti-protease activity, at current therapeutic dosages its activity enhances the plasmatic concentration of the main antiviral compound by its ability to block drug metabolism. However, in an open-label clinical trial using LPV/RTV against COVID-19, their combination showed a limited benefit for treated patients^14^. In the early 2000s, another contemporary antiretroviral protease inhibitor, atazanavir (ATV), replaced LPV/RTV due to fewer side effects for the patients^15,16^. Contemporarily, *in silico* evidence suggested that other HIV protease inhibitors would target SARS-CoV-2 Mpro better than LPV or RTV, that included ATV^17^. Importantly, ATV has been described to reach the lungs after intravenous administration^18,19^. Moreover, a proposed secondary use of ATV to treat pulmonary fibrosis suggested that this drug could functionally reach the lungs^19^.

The seriousness of COVID19 and the need for an immediate intervention, along with this series of observations with LPV, RTV and ATV, motivated us to evaluate the susceptibility of SARS-CoV-2 to ATV. Since ATV is available as a clinical treatment alone or in combination with RTV, both therapies were studied here, which for the first time describes that SARS-CoV-2 Mpro is a target for ATV. Further, ATV alone or withRTV could inhibit viral replication in cell culture models of infection that also prevented the release of a cytokine storm-associated mediators. Our timely data highlights an additional therapeutic approach against COVID-19 that should be considered for clinical trials.

## 2) Results

### 2.1) ATV docks into SARS-CoV-2 Mpro more spontaneously and stably than LPV

The targeting of the enzyme Mpro from SARS-CoV-2 by both ATV and LPV was evaluated by molecular modeling using a representative structure (PDB:6LU7). As shown in Figure 1, ATV occupied the S1* and S1 regions, whereas LPV occupied S1* and S2 regions with calculated free energy scores for LPV and ATV of −59.87 and - 65.49 Kcal/mol, respectively. The more spontaneous binding of ATV, suggested by its lower energy score, may be related to its projected ability to form hydrogens bonds with the amino acid residues Asn142, His164, and Glu166 in Mpro, whereas the binding of LPV depends on hydrophobic interactions (Figure 2).

**Figure 1.**
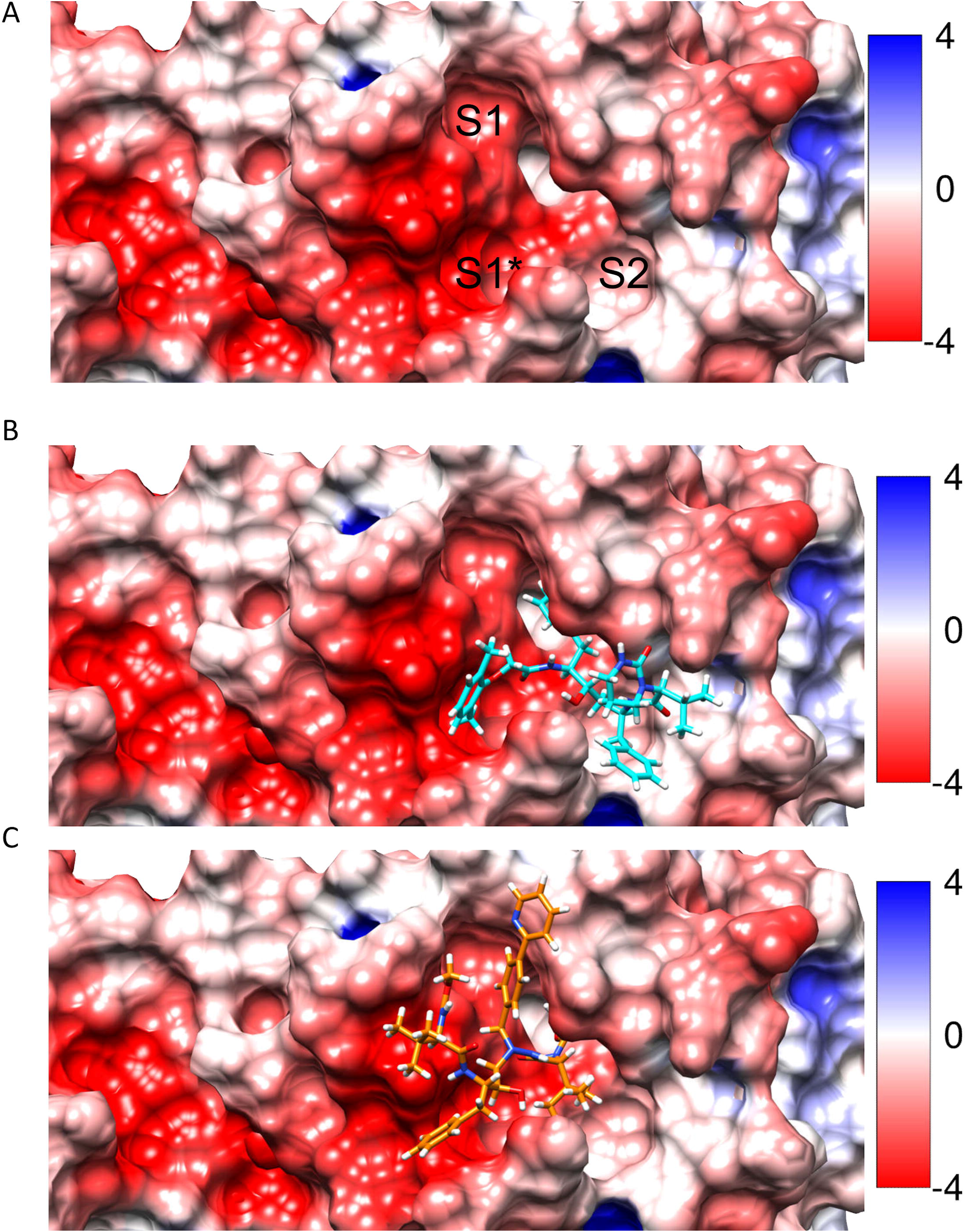
The active site of SARS-CoV-2 Mpro in the absence and presence of the inhibitors. A representative structure of Mpro (PDB:6LU7) was color coded to show the electrostatic potential of residues in the active site for negative (blue) and positive (red) charges. Panel A, the cavities of ligand interaction designated S1*, S1 and S2 in the absence of inhibitors. Panel B, placement of LPV (cyan) docked in the S1* and S2 regions of the active site. Panel C, placement of ATV (orange) docked in the S1* and S1 regions of the active site.

**Figure 2.**
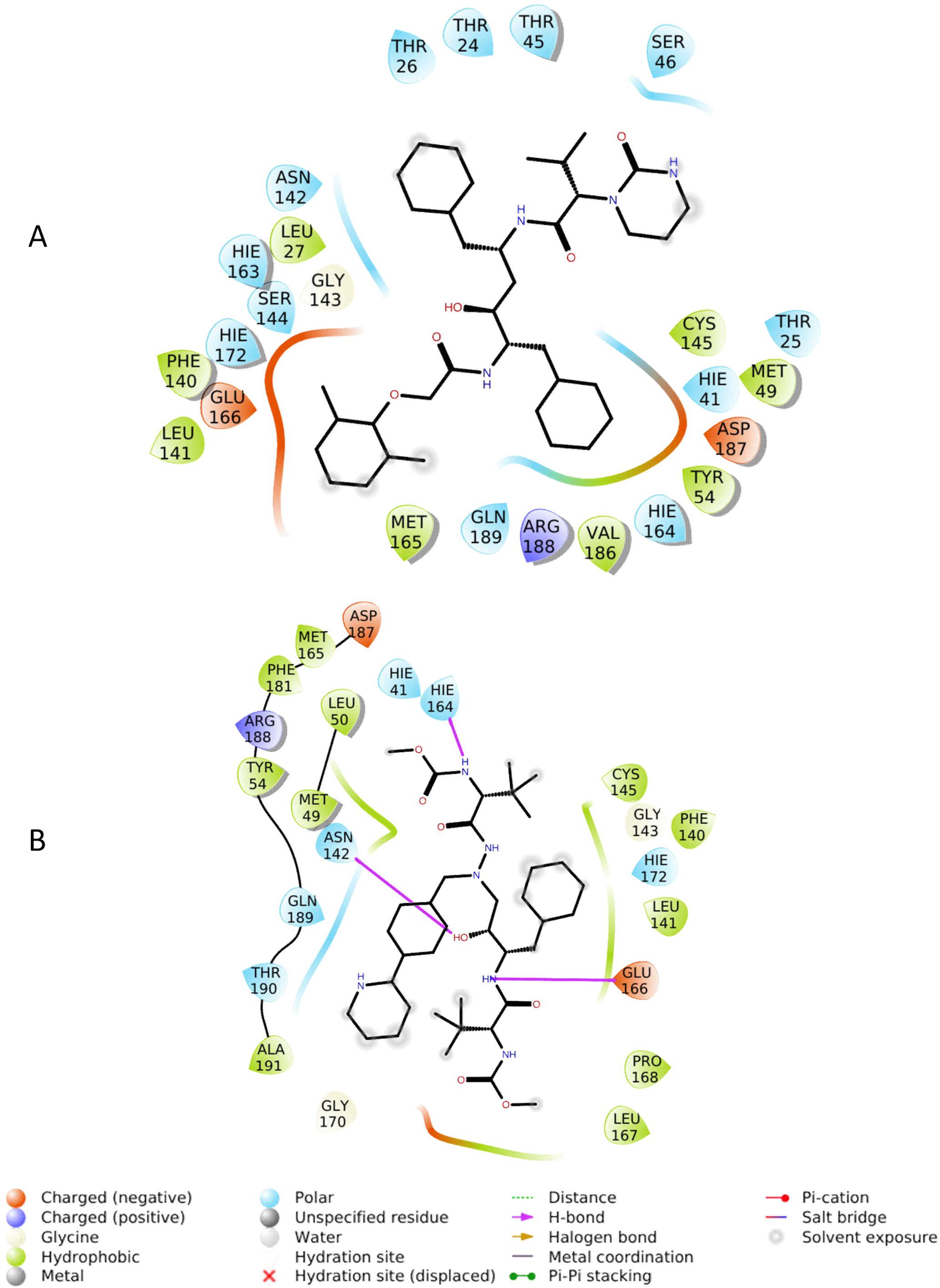
Binding profile of antiretroviral drugs onto SARS-CoV-2 Mpro. Two-dimensional (2D) representations of the interactions of LPV (A) and ATV (B) in the Mpro active site based on a molecular docking analysis. Two hydrogen bonds are predicted between ATV and Mpro.

A molecular dynamic analysis revealed that the root-mean-square deviation (RMSD) for the SARS-CoV-2 Mpro backbone presented different conformations in complex with ATV or LPV (Figure S1). LPV was initially at a 3.8 Å distance from the catalytic residue Cys145 (Figure S2A and S3A), which after conformational changes extended to a distance equivalent to 7.17 Å (Figure 3A and 4A) that is projected to most likely limit the extent of its antiviral inhibition. Another critical residue, His41, was satisfactorily at a distance of 2.89 Å from bound LPV (Figure 3A and 4A). While ATV did not interact with His41 or Cys145 (Figure S2B and S3B), its position remained stable within the active site independent of conformational changes displayed by the enzyme (Figure 3B and 4B). The steric occupation of the cleft in the enzymatic active site by ATV, which block the residues of the catalytic amino acids, can be explained by its stronger interactions with Mpro, compared to LPV, through multiple hydrogen bonds during stationary docking and molecular dynamics (Tables S1-S3).

**Figure 3.**
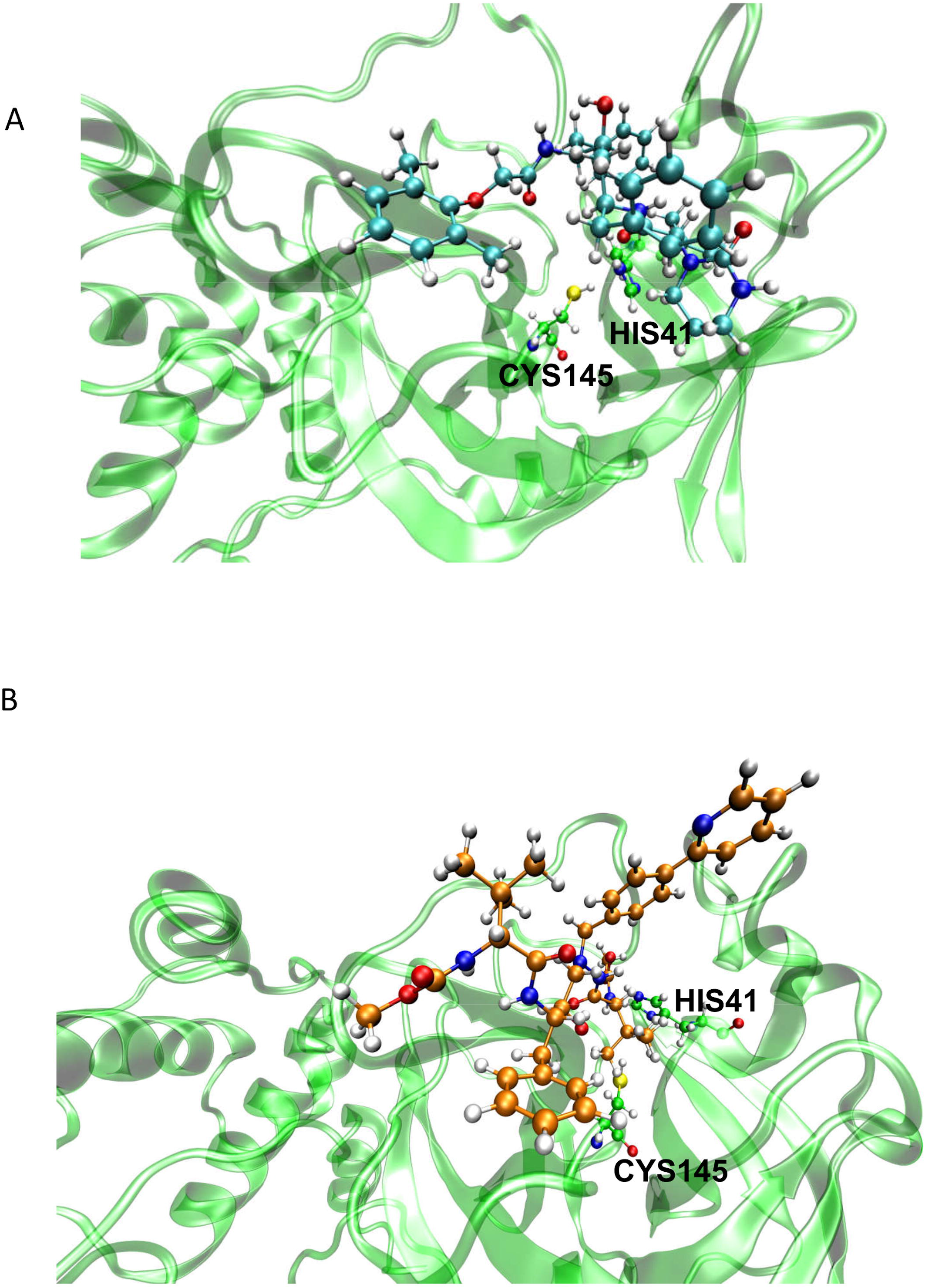
Final positions of ATV and LPV on Mpro at the end of a molecular dynamic simulation. Representative images of the molecular dynamics after 100 ns of simulation. LPV (A) and ATV (B) are positioned in the Mpro active site at the end of 100 ns simulation.

**Figure 4.**
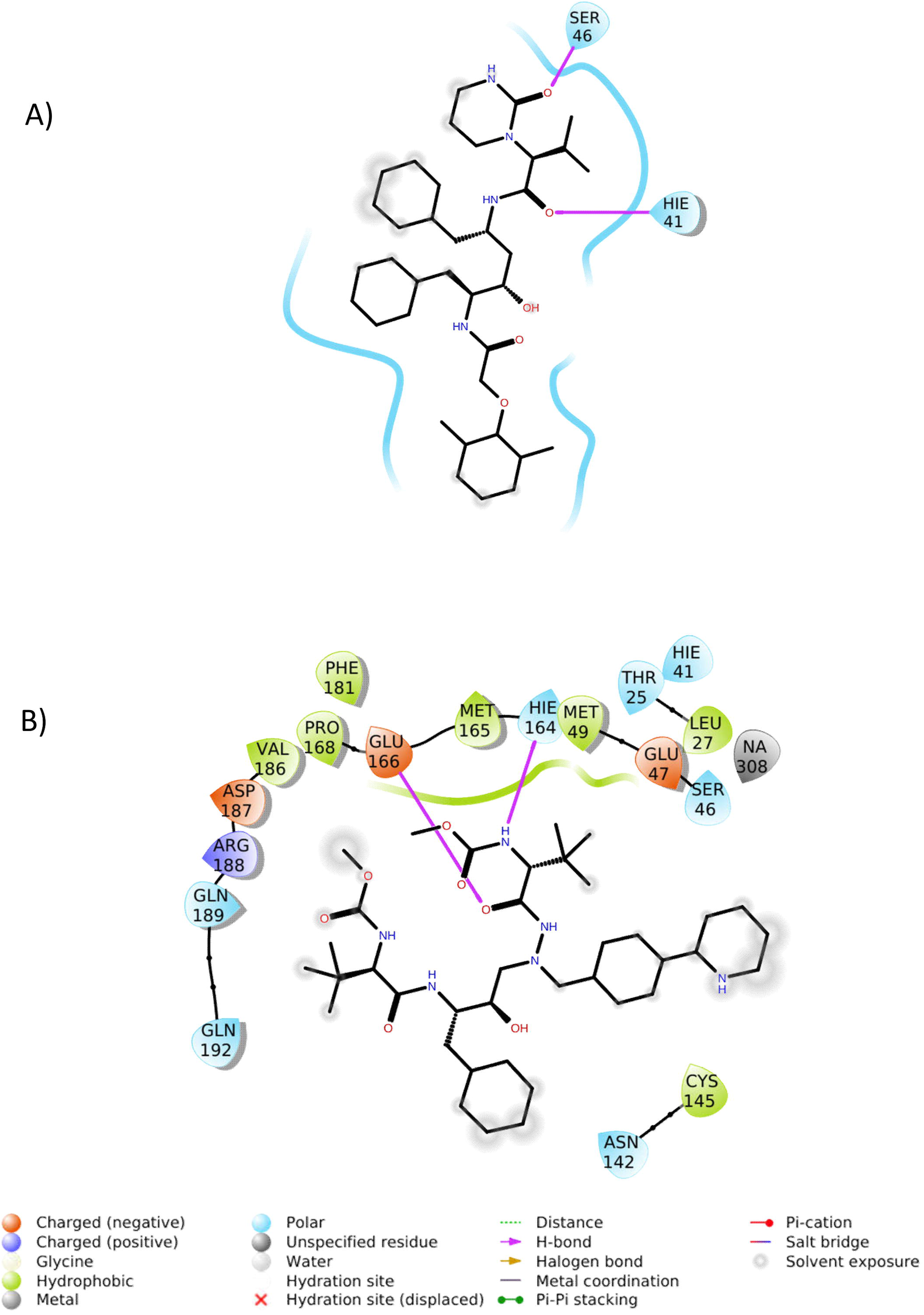
Position profile of ATV and LPV during molecular dynamics. Two-dimensional (2D) representation of the interactions of LPV (A) and ATV (B) in the Mpro active site at the end of 100 ns molecular dynamic simulation.

### 2.2) ATV inhibits SARS-CoV-2 Mpro enzymatic activity

Next, we evaluated whether ATV could inhibit SARS-CoV-2 Mpro activity by partially purifying the enzyme in cellular fractions obtained from SARS-CoV-2-infected cells and performing zymographic profiles. To assure that the proteinase profiles were not dependent on cellular enzymes, similar fractions of mock-infected cells were also prepared for comparison. The results from cysteine proteinase zymographic profiles in gelatinolytic gels reveled a cellular related band of approximately 70 kDa under both conditions (Figure 5, lanes Nil). This activity was blocked by the drug E-64, an epoxide that acts as an irreversible inhibitor of cysteine proteases (Figure 5, lanes E-64). In the infected cells, a region of activity was observed between 31 and 38 kDa that was not present in the mock fraction. This zone of molecular weight is consistent with expected size of SARS-CoV-2 Mpro as was the inhibition of activity in this region by exposure of the gels to 10 μM of ATV, which did not affect the cellular cysteine proteinase at 70 kDa (Figure 5, lanes ATV). Further confirmation of the presence and activity of SARS-CoV-2 Mpro in fractions from infected cells was obtained by treatment with RTV, which inhibited activity in the molecular range of 31-38 kDa without a change in the 70 kDa region (Figure 5, lanes RTV). These data are consistent with predictions from the molecular modeling and dynamic analyses that suggested that ATV could bind and target the enzymatic activity of the Mpro encode by the novel 2019 CoV.

**Figure 5.**
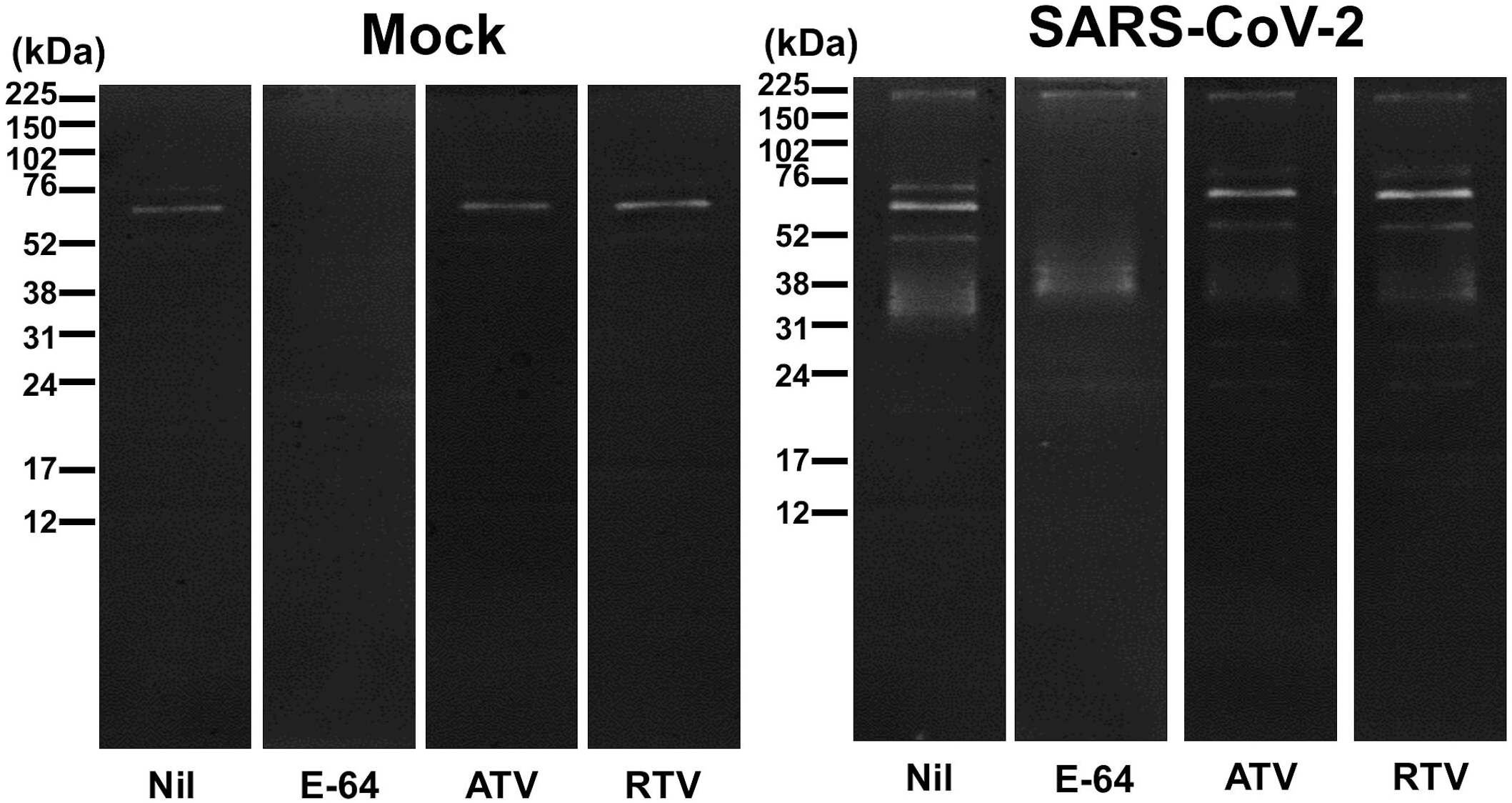
Inhibition of proteinase activity through an analysis of gelatinolytic activity. Vero cells were mock treated or infected with SARS-CoV-2 at an MOI of 0.1 for 48h before lysis and preparation of a cellular fraction. Fractions containing 12 µg of total protein separated by electrophoresis followed by cutting the gels into their individual lanes that were incubated in 10 mM sodium acetate buffer (pH 5.5) in the absence (Nil) or presence of 10 µM of E-64, ATV or RTV. Gelatinolytic bands indicative of enzymatic activity were revealed by negative staining with amide black solution. Molecular mass markers are indicated (kDa).

### 2.3) SARS-CoV-2 is susceptible to ATV in different cell types

We extended our investigation to the inhibition of SARS-CoV-2 replication by ATV using a range of different cellular systems. Vero cells are a well-known model system that produce high virus titers and display visual cytopathic effects to viral infections. ATV alone, or in combination with RTV, inhibited infectious virus production and SARS-CoV RNA levels in Vero cells (Figure 6A and B, respectively). CQ was used as a positive control because of its inclusion in the SOLIDARITY trial due to its encouraging pre-clinical and clinical results against SARS-CoV-2 replication and COVID-19, respectively^20,21^. ATV/RTV was the most potent therapy tested; with an EC_50_ of 0.5 ± 0.08 µM. ATV alone and CQ’s potencies were 2.0 ± 0.12 µM and 1.0 ± 0.07 µM, respectively. SARS-CoV-2 susceptibility to CQ is consistent with recent reports in the literature^20^, validating our analysis. The ATV/RTV, ATV, and CQ cytotoxicity values, CC_50_, were 280 ± 3 µM, 312 ± 8 µM and 259 ± 5 µM, respectively. Our results indicate that the selectivity index (SI, which represents the ratio between the CC_50_ and EC_50_ values) for ATV/RTV, ATV and CQ were 560, 156 and 259, respectively, which shows that ATV/RTV has a high therapeutic potential that was greater than CQ.

**Figure 6.**
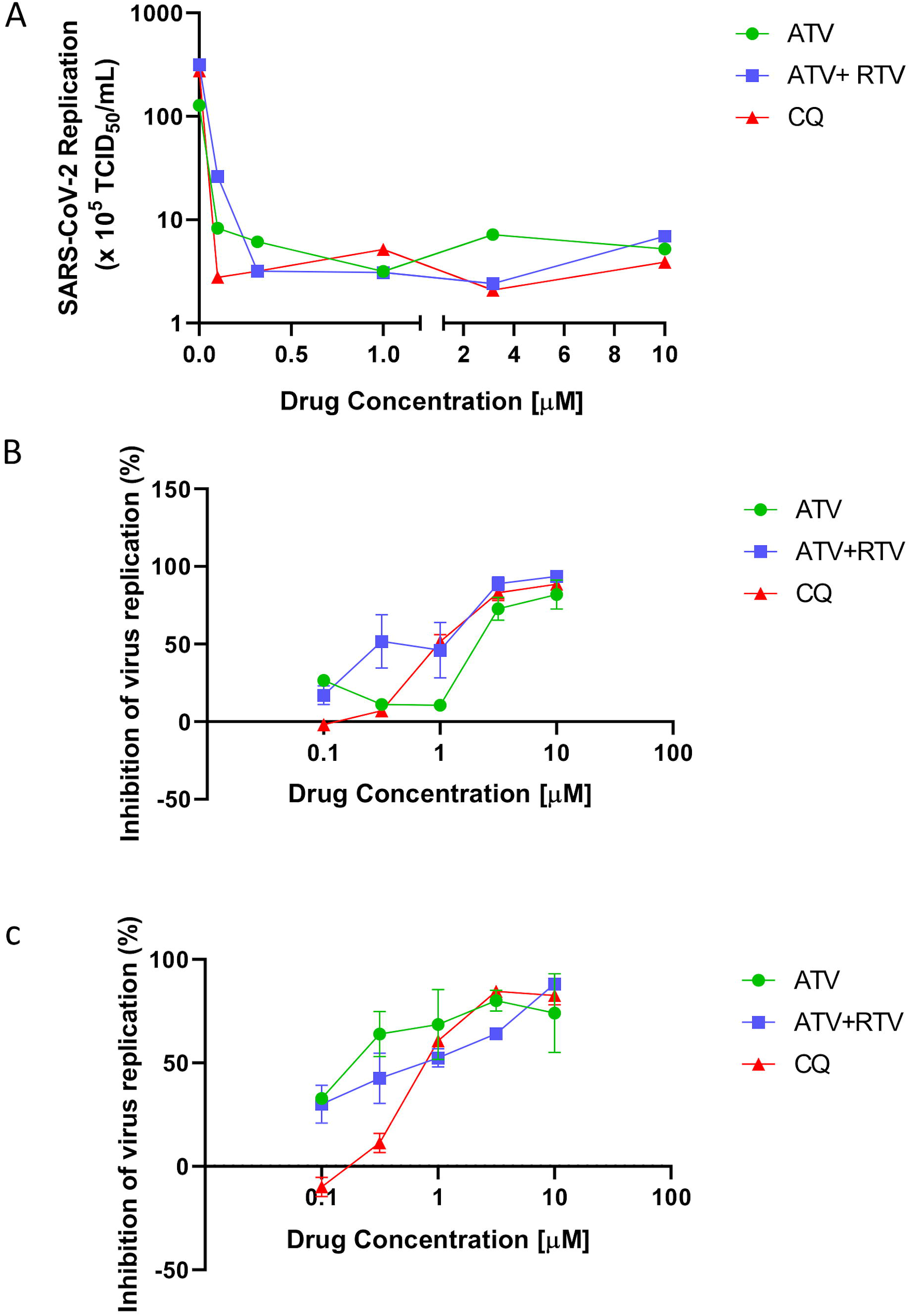
The antiviral activity of ATV and ATV/RTV against SARS-CoV-2. Vero (A and B) or A549 (C) cells were infected with SARS-CoV-2 at the MOI of 0.01 and exposed to indicated concentrations of the drugs. After 2 days, the viral replication in the culture supernatant was measured by TCID_50_/mL (A) or RT-PCR (B and C). The data represent means ± SEM of three independent experiments.

Since the results regarding the pharmacologic activity of ATV and ATV/RTV against SARS-CoV-2 replication in Vero cells were promising, we next investigated whether the proposed drug therapies could inhibit virus replication in a human epithelial pulmonary cell line (A549). ATV alone showed a nearly 10-fold increase in potency for inhibiting SARS-CoV-2 replication in A549 (Figure 6C) compared to Vero cells (Figure 6B). ATV/RTV and CQ were similarly potent in inhibiting virus replication in both cell types (Figure 6B and C). ATV/RTV, ATV and CQ EC_50_ values to inhibit SARS-CoV-2 replication in A549 cells were 0.60 ± 0.05 µM, 0.22 ± 0.02 µM and 0.89 ± 0.02 µM, respectively. *In vitro* results confirmed the rational that SARS-CoV-2 would be susceptible to ATV that included cells derived from the respiratory tract.

### 2.4) ATV prevents cell death and pro-inflammatory cytokine production in SARS-CoV-2-infected monocytes

Recent reports on the COVID-19 outbreak have implicated that an increase in the levels of lactate dehydrogenase (LDH) and interleukin 6 (IL-6) is associated with mortality^22^. Viral infection in the respiratory tract often trigger the migration of blood monocytes to orchestrate the transition from innate to adaptive immune responses^23^. For these reasons, ATV and ATV/RTV were tested at suboptimal (1 μM) or optimal (10 μM) doses in a SARS-CoV-2-infection model utilizing human primary monocytes.

ATV/RTV and CQ were similarly efficient to inhibit viral replication in the human monocytes (Figure 7A). Virus infection increased cellular mortality by 75%, which was prevented by ATV, at both doses tested, and by ATV/RTV, at 10 μM (Figure 7B). As a control, detergent treatment completely destroyed all cells (Figure 7B). Moreover, we observed that infections by SARS-CoV-2 triggered the expected increase in the IL-6 levels in the culture supernatant, which ranged from 20- to 60-fold depending on the cell donor (Figure 7C, open circles in nil-treated cells). The virus-induced enhancement of IL-6 levels were significantly prevented by treatment with ATV at 10 µM, ATV/RTV at both 1 and 10 µM and CQ at 10 µM (Figure 7C). Another biomarker of uncontrolled pro-inflammatory cytokine response, TNF-α, was up-regulated 40-fold during virus infection (Figure 7D). Only the combination of ATV/RTV could significantly prevent the induction of TNF-α release (Figure 7D). Altogether, our results confirm that ATV and ATV/RTV should not be ignored as an additional therapeutic option against COVID-19.

**Figure 7.**
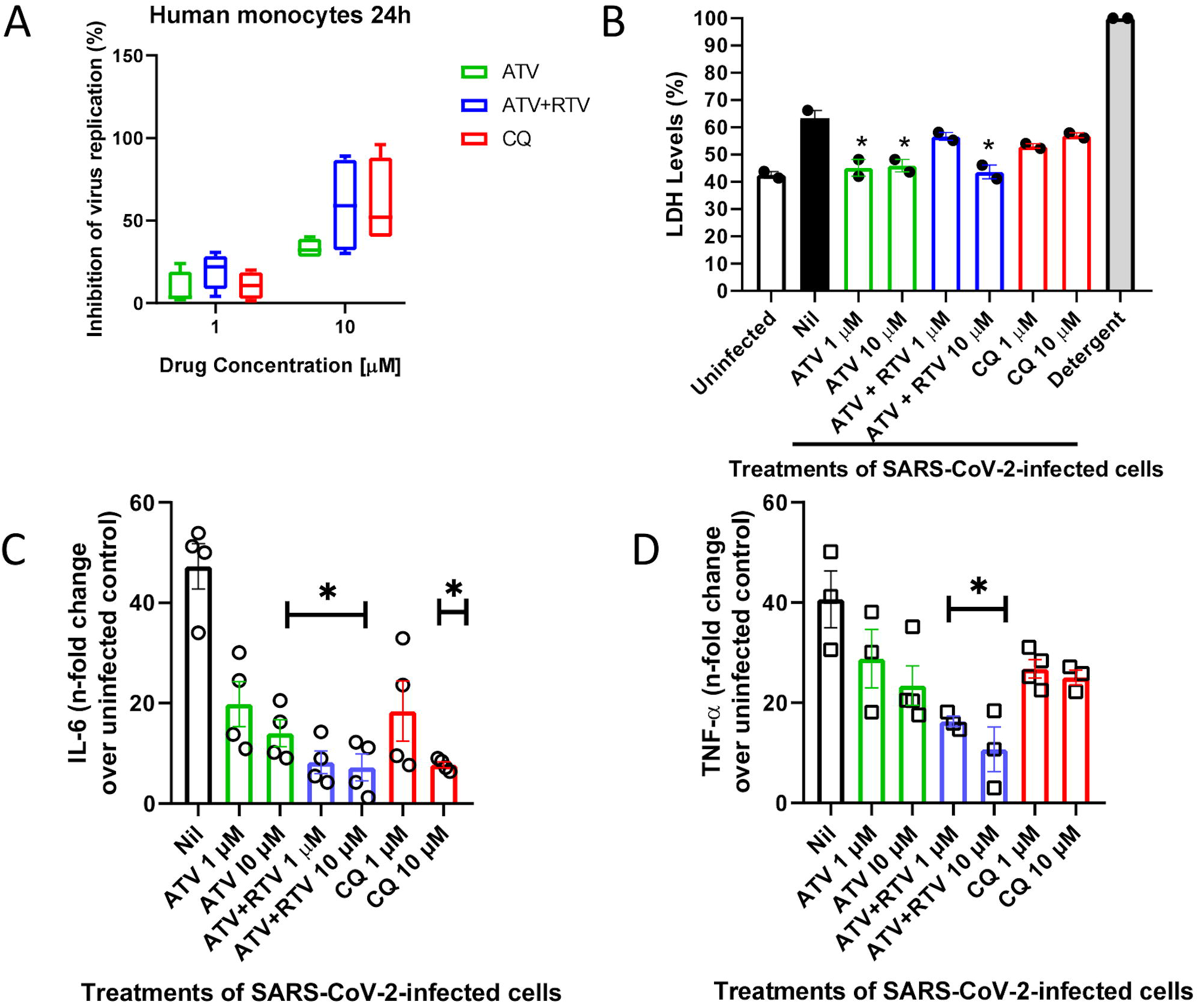
ATV and ATV/RTV impairs SARS-CoV-2 replication, cell death and cytokine storm in human primary monocytes. Human primary monocytes were infected at the indicated MOI of 0.01 and treated with indicated concentration of the compounds. After 24h, virus replication (A) and LDH release (B) as well as the levels of IL-6 (C) and TNF-α (D) were measured in the culture supernatant. The data represent means ± SEM of experiments with cells from at least three healthy donors. Differences with *P <* 0.05 are indicates (*), when compared to untreated cells (nil).

## 3) Discussion

In these two decades of the 21st century, the human vulnerability to emerging viral diseases has been notable^24^. The emergence of infectious disease highlights the undeniable fact that existing countermeasures are inefficient to prevent virus spill over and diseases outbreak. Preclinical data on the susceptibility of an emerging virus to clinically approved drugs can allow for the rapid mobilization of resources towards clinical trials^8^. This approach proved feasible for combating the Zika, yellow fever and chikungunya outbreaks experienced in Brazil over the past 5 years, when our group demonstrated that sofosbuvir, a blockbuster drug against hepatitis C, could represent a compassionate countermeasure against these diseases^25–29^.

Currently, the rate of SARS-CoV-2 dissemination has become one of the most rapidly evolving pandemics known in modern times with the number of cases and deaths doubling every week and the peak of the pandemic has yet to arrive^6^. The existence of several ongoing clinical trials against COVID-19 reinforces the suggestion that drug repurposing represents the fastest approach to identify therapies to emerging infectious disease^8^. The WHO/UN, under the auspicious of the SOLIDARITY trial, have highlighted the most promising anti-CoV drugs, such as LPV/RTV with or without interferon-β, CQ and remdesivir^30^. Here, we provide preclinical evidence that another HIV protease inhibitor, ATV, can inhibit the activity of a critical protease of SARS-CoV-2, Mpro, and that this inhibition can extend to a disruption of viral replication as well as the release of cytokine storm-associated mediators associated with viral infection. The results suggest that the performance of ATV could be better than LPV and strongly support that inclusion of ATV-based therapies in clinical trials for COVID-19 either alone, in combination with RTV or both.

Kaletra® is an LPV/RTV formulation from Abbot Laboratories that was approved by Food and Drug Administration (FDA) in 2000 and was evaluated for use in the treatment during the SARS-CoV outbreak in 2002 and again for MERS-CoV. Its continued evaluation with the SARS-CoV-2 outbreak is a logical choice due to the conservation of the Mpro among these highly pathogenic viruses^31,32^. Nevertheless, information on the susceptibility of SARS-CoV-2 to other antiviral protease inhibitors that have been approved since 2003 has been scarce.

ATV was approved in 2003, and become a wider prescribed drug among HIV-infected individuals, than LPV, including for critically ill patients^16^. ATV shows a safer profile than LPV in both short- and long-tem therapeutic regimens^15,33^. ATV has a documented bioavailability to reach the respiratory tract^18,34^, which lead to its proposed use against pulmonary fibrosis^19^. However, it is not currently under consideration for clinical trials against COVID-19.

The potencies of LPV and LPV/RTV against CoV are from 10 to 8 μM, respectively^32^. Based on our data, ATV and ATV/RTV are at least 10 times more potent. The ATV and ATV/RTV *in vitro* potencies are comparable to other small molecule inhibitors of the SARS-CoV-2, such as remdesivir and CQ^20^. The improved potency of ATV, in comparison to LPV, may be at least in part due to its multiple hydrogen bond driven interactions within the Mpro active site. Other investigators have also recognized a wider range of interactions of ATV and Mpro compared to LPV^17,35^, although none provided functional evidence through phenotypic assays as presented here. Neither ATV nor LPV displayed any interactions with the catalytic dyad of Cys145 and His41 at the start of the molecular dynamic simulations. However, important interactions were observed at its end, such as LPV-His41 and ATV-Glu166. Glu166 is one of the residues that promotes the opeing of Mpro for its substrate to interact with the active site^36,37^.

LPV/RTV was the first line of defense for early patients with COVID-19^31^. In patients with severe COVID-19, the open-labeled clinical trial with LPV/RTV revealed that treated patients had 5% less deaths and better clinical improvement then controls^14^. Throughout the course of this study, LPV/RTV-treated patients continued to shed SARS-CoV-2 at the same magnitude and duration of the control group^14^, limiting the enthusiasm on the part of the medical and scientific community for this therapeutic option. In this context, ATV and/or ATV/RTV should not be ignored in the treatment of this novel respiratory disease. Indeed, our results demonstrate that the potency of ATV and ATV/RTV potency against SARS-CoV-2 in A549 cells is likely to be consistent with their bioavailability in the lungs in experimental models^18,34^.

Highly pathogenic respiratory viruses, such as influenza A virus, have been associated with a cytokine storm that describes an uncontrolled pro-inflammatory cytokine response^38,39^. Cytokine storms also seem to be highly relevant for pathogenic human CoVs^40^. Contemporary investigations on SARS-CoV-2 strongly suggest the involvement of cytokine storm with disease severity^22^. COVID-19 mortality is associated with enhanced IL-6 levels and consistent cell death, as measured by LDH release^22^. We showed that ATV and ATV/RTV decreased IL-6 release in SARS-CoV-2-infected human primary monocytes. Moreover, we also included in our analysis TNF-α, another hallmark of inflammation during respiratory virus infections^22,41^. Our results reveled that cellular mortality and cytokine storm-associated mediators were reduced after treatment with the repurposed antiretroviral drugs used in this study.

Among the most promising anti-SARS-CoV-2 drugs, CQ, IFN-β and LPV displayed a higher toxic profile than ATV. Moreover, ATV and ATV/RTV have *in vitro* antiviral potencies comparable to CQ and remdesivir, which were superior to LPV/RTV. In summary, our study highlights a new option among clinically approved drugs that should be considered in ongoing clinical trials for an effective treatment for COVID-19.

## Material and Methods

### 4.1. Reagents

The antiviral ATV, ATV/RTV and CQ were received as donations from Instituto de Tecnologia de Fármacos (Farmanguinhos, Fiocruz). ATV/RTV was prepared in the proportion of 3:1 as the pharmaceutical pills are composed of 300 mg ATV and 100 mg RTV daily. ELISA assays were purchased from R&D Bioscience. All small molecule inhibitors were dissolved in 100% dimethylsulfoxide (DMSO) and subsequently diluted at least 10^4^-fold in culture or reaction medium before each assay. The final DMSO concentrations showed no cytotoxicity. The materials for cell culture were purchased from Thermo Scientific Life Sciences (Grand Island, NY), unless otherwise mentioned.

Triton X-100 (TX-100), 3-[(3-Cholamidopropyl)dimethylammonio]-1-propanesulfonate hydrate (CHAPS), 1,2,3-Propanetriol (glycerol), bovine serum albumin (BSA), Phosphate-buffered saline (PBS), N-benzyloxycarbonyl-l-phenylalanyl-l-arginine 7-amino-4-methylcoumarin (Z-FR-AMC; ε= 1.78 × 10^4^ M^−1^ cm^−1^), dithiothreitol (DTT) and trans-epoxysuccinyl-l-leucylamido(4-guanidino)butane (E-64) were purchased from Sigma Aldrich Chemical Co. (St. Louis, MO, USA). HiTrap Q FF anion exchange chromatography column (HiTrap Q FF) was purchase from GE Healthcare Life Sciences. Micro-bicinchoninic acid (BCA) protein assay kit was purchased from Pierce Chemical Co. (Appleton, WI). All other reagents were of analytical grade or better.

### 4.2. Cells and Virus

African green monkey kidney (Vero, subtype E6) and A549 (human lung epithelial cells) cells were cultured in high glucose DMEM with 10% fetal bovine serum (FBS; HyClone, Logan, Utah), 100 U/mL penicillin and 100 μg/mL streptomycin (Pen/Strep; ThermoFisher) at 37 °C in a humidified atmosphere with 5% CO_2_.

Human primary monocytes were obtained after 3 h of plastic adherence of peripheral blood mononuclear cells (PBMCs). PBMCs were isolated from healthy donors by density gradient centrifugation (Ficoll-Paque, GE Healthcare). PBMCs (2.0 × 10^6^ cells) were plated onto 48-well plates (NalgeNunc) in RPMI-1640 without serum for 2 to 4 h. Non-adherent cells were removed and the remaining monocytes were maintained in DMEM with 5% human serum (HS; Millipore) and penicillin/streptomycin. The purity of human monocytes was above 95%, as determined by flow cytometric analysis (FACScan; Becton Dickinson) using anti-CD3 (BD Biosciences) and anti-CD16 (Southern Biotech) monoclonal antibodies.

SARS-CoV-2 was prepared in Vero E6 cells from an isolate contained on a nasopharyngeal swab obtained from a confirmed case in Rio de Janeiro, Brazil. Viral experiments were performed after a single passage in a cell culture in a 150 cm^2^ flasks with DMEM plus 2% FBS. Observations for cytopathic effects were performed daily and peaked 4 to 5 days after infection. All procedures related to virus culture were handled in a biosafety level 3 (BSL3) multiuser facility according to WHO guidelines. Virus titers were determined as the tissue culture infectious dose at 50% (TCID_50_/mL). Virus stocks were kept in - 80 °C ultralow freezers.

The virus strain was sequenced to confirm the virus identity and its complete genome is publicly deposited (https://nextstrain.org/ncov: Brazil/RJ-314/2020 or GISAID EPI ISL #414045).

### 4.3. Cytotoxicity assay

Monolayers of 1.5 × 10^4^ Vero cells in 96-well plates were treated for 3 days with various concentrations (semi-log dilutions from 1000 to 10 µM) of ATV, ATV/RTV or CQ. Then, 5 mg/ml 2,3-bis-(2-methoxy-4-nitro-5-sulfophenyl)-2*H*-tetrazolium-5-carboxanilide (XTT) in DMEM was added to the cells in the presence of 0.01% of N-methyl dibenzopyrazine methyl sulfate (PMS). After incubating for 4 h at 37 °C, the plates were measured in a spectrophotometer at 492 nm and 620 nm. The 50% cytotoxic concentration (CC_50_) was calculated by a non-linear regression analysis of the dose– response curves.

### 4.4. Yield-reduction assay

Cells were infected with a multiplicity of infection (MOI) of 0.01. Vero or A549 cells were infected at densities of 5 × 10^5^ cells/well. Human primary monocytes were infected at density of 2-8 × 10^5^ cells/well, depending on the endogenous characteristic of the cell donor. Infections were performed in 48-well plates for 2h at 37 °C. The cells were washed, and various concentrations of compounds were added to DMEM with 2% FBS. After 48h, supernatants were collected and harvested virus was quantified by real time RT-PCR and infectious titers by TCID_50_/mL. A variable slope non-linear regression analysis of the dose-response curves was performed to calculate the concentration at which each drug inhibited the virus production by 50% (EC_50_).

### 4.5. Virus titration

Monolayers of Vero cells (2 × 10^4^ cell/well) in 96-well plates were infected with a log-based dilution of supernatants containing SARS-CoV-2 for 1h at 37°C. Cells were washed, fresh medium added with 2% FBS and 3 to 5 days post infection the cytopathic effect was scored in at least 10 replicates per dilution by independent readers. The reader was blind with respect to source of the supernatant. A Reed and Muench scoring method was employed to determine TCID_50_/mL^42^.

### 4.6. Molecular detection of virus RNA levels

The total RNA from a culture was extracted using QIAamp Viral RNA (Qiagen®), according to manufacturer’s instructions. Quantitative RT-PCR was performed using QuantiTect Probe RT-PCR Kit (Quiagen®) in an ABI PRISM 7500 Sequence Detection System (Applied Biosystems). Amplifications were carried out in 25 µL reaction mixtures containing 2× reaction mix buffer, 50 µM of each primer, 10 µM of probe, and 5 µL of RNA template. Primers, probes, and cycling conditions recommended by the Centers for Disease Control and Prevention (CDC) protocol were used to detect the SARS-CoV-2^43^. The standard curve method was employed for virus quantification. For reference to the cell amounts used, the housekeeping gene RNAse P was amplified. The Ct values for this target were compared to those obtained to different cell amounts, 10^7^ to 10^2^, for calibration.

### 4.7. Measurements Inflammatory Mediators and cell death marker

The levels of TNF-α, IL-6 and LDH were quantified in the monocyte supernatants from infected and uninfected cells. ELISA for TNF-α and IL-6 required 100 µL of supernatants to be exposed to capture antibody in 96-well plates. After a 2h incubation period at room temperature (RT), the detection antibody was added. Plates were incubated for another 2h at RT. Streptavidin-HRP and its substrate were added, incubated for 20 minutes and the optical density was determined using a microplate reader set to 450 nm.

Extracellular lactate dehydrogenase (LDH) was quantified using Doles® kit according to manufacturer’s instructions. Supernatant was centrifuged at 5,000 rpm for 1 minute, to remove cellular debris. A total of 25 µL of supernatant was placed into 96-well plates and incubated with 5 µL of ferric alum and 100 µL of LDH substrate for 3 minutes at 37 °C. Nicotinamide adenine dinucleotide (NAD, oxidized form) was added followed by the addition of a stabilizing solution. After a 10 min incubation, plates were measured in a spectrophotometer at 492 nm.

### 4.8. Molecular docking

ATV (PubChem CID: 148192) and LPV (PubChem CID: 92727) were used as inhibitors of the SARS-CoV-2 Mpro. ATV and LPV were prepared using the Generalized Amber Force Field (GAFF) and their charges were obtained using the AM1-BCC loading scheme ^44,45^.

Molecular docking experiments were performed with DOCK 6.9^46^ for identifying the binding site of the Mpro. SARS-CoV-2 Mpro structure was obtained from Protein Data Bank (RCSB PDB, http://www.rcsb.org), under the accession code #6LU7 ^47^. The active site region was identified by using a complexed peptide (N-[(5-methylisoxazol-3-yl)carbonyl]alanyl-l-valyl-n∼1∼-((1r,2z)-4-(benzyloxy)-4-oxo-1-{[(3r)-2-oxopyrrolidin-3-yl]methyl}but-2-enyl)-l-leucinamide) as a guide. The creation of the DOCK 6.9 input files for docking was performed using Chimera 1.14^48^.

The docking of ligands was performed in a box of 10 Å edges with its mass center matching that of the complexed peptide. Each scan produced 20 conformations for each ligand with the best score being used for molecular dynamics simulations.

### 4.9. Molecular dynamics

Since the tertiary structure (3D) of the SARS-CoV-2 Mpro is a homodimer, we focused the molecular dynamics only one chain, henceforward chain A. Molecular dynamics calculations were performed using NAMD 2.9^49^ and Charmm27* force field^50^ at pH 7, i.e., with deprotonated Glu and Asp, protonated Arg and Lys, and neutral His with a protonated Nε atom. This all-atom force field has been able to fold properly many soluble proteins^51–53^. The soluble proteins were centered in a cubic box of TIP3P water molecules^54^; the box extended 1.2 nm outside the protein on its four lateral sides, and the appropriate numbers of Na+ and Cl-ions were added to ensure system neutralization. The electrostatic interactions were calculated using the Particle Mesh Ewald method and a cutoff of 1.2 nm^55^. The same cutoff of 1.2 nm was used for the Van der Waals interactions. The non-bonded pair lists were updated every 10 fs. In what follows, the analysis is based on MD simulation of 100 ns at 310 K.

### 4.10. Protein extraction

Protein extracts containing SARS-CoV-2 Mpro activity were obtained from Vero cell monolayers at 25 cm^2^ flasks that were infected for 1h with an MOI of 0.1 at 37 °C and 5% CO_2_. After 1 or 2 days of infection, the supernatant was harvested and monolayers were washed 3 times with in sterile cold PBS (pH 7.2). Next, cells were suspended into 1 mL of lysis buffer (100 mM Tris-HCl (pH 8.0), 150 mM NaCl, 10% glycerol and 0.6% Triton X-100) and kept at 4 °C. The soluble protein fraction was isolated as the supernatant after centrifugation (100,000 x g, 30 min, 4 °C) and stored at −20°C until further use. The protein concentrations of the samples were determined using the BCA protein assay kit.

### 4.11. Zymographic assays

Proteinases were assayed after electrophoresis on 10% SDS-PAGE with 0.1% copolymerized gelatin^56^. Briefly, the gels were loaded per slot with 12 μg of soluble proteins dissolved in Laemmli’s buffer, and following electrophoresis at a constant voltage of 200 V at 4°C, they were soaked for 1 h at 25 °C in washing buffer (0.1 mM sodium acetate buffer (pH 5.5) containing 2.5% TX-100). Proteinase activity was detected by incubating (16 h at 37 °C) the gels in reaction buffer (0.1 mM sodium acetate buffer pH 5.5 containing 1.0 mM DTT), in the presence and absence of same concentration of 10 µM of E-64, ATV, RTV or the ATV/RTV combination. Hydrolysis of gelatin was visualized by staining the gels with amido black 0.2%^57^.

### 4.12. Statistical analysis

The assays were performed blinded by one professional, codified and then read by another professional. All experiments were carried out at least three independent times, including a minimum of two technical replicates in each assay. The dose-response curves used to calculate EC_50_ and CC_50_ values were generated by variable slope plot from Prism GraphPad software 8.0. The equations to fit the best curve were generated based on R^2^ values ≥ 0.9. Student’s T-test was used to access statistically significant *P* values <0.05. The statistical analyses specific to each software program used in the bioinformatics analysis are described above.

## Author contributions

Experimental execution and analysis – NFR, CQS, CRL, FSS, ACF, MM, ARM, CSF, VCS, SSGD, MM, AM

Data analysis, manuscript preparation and revision – NFR, CQS, ACF, CSF, CRL, FSS, CRA, FAB, MMS, PTB, TMLS

Conceptualized the experiments – NFR, CQS, TMLS Study coordination – TMLS

Manuscript preparation and revision – PTB, TMLS

## Acknowledgments

Thanks are due to Dr. Carmen Beatriz Wagner Giacoia Gripp for assessments related to BSL3 facility. This work was supported by Conselho Nacional de Desenvolvimento Científico e Tecnológico (CNPq), Fundação de Amparo à Pesquisa do Estado do Rio de Janeiro (FAPERJ). This study was financed in part by the Coordenac□ão de Aperfeic□oamento de Pessoal de Nível Superior - Brasil (CAPES) - Finance Code 001. Funding was also provided by CNPq, CAPES and FAPERJ through the National Institutes of Science and Technology Program (INCT) to Carlos Morel (INCT-IDPN). Thanks are due to Oswaldo Cruz Foundation/FIOCRUZ under the auspicious of Inova program. The funding sponsors had no role in the design of the study; in the collection, analyses, or interpretation of data; in the writing of the manuscript, and in the decision to publish the results.

**The authors declare no competing financial interests.**

