## Supplementary material for "Atazanavir inhibits SARS-CoV-2 replication and pro-inflammatory cytokine production": Figures S1-S3 and Tables S1-S3 (from another version of this preprint: version 1, 2020-04-05)

2 – Laboratório de Biologia Molecular e Doenças Endêmicas, IOC, Fiocruz, Rio de Janeiro, RJ, Brazil.

3 – Universidade Iguaçu, Nova Iguaçu, RJ, Brazil.

4 – Laboratório de Pesquisas em Timo, IOC, Fiocruz, Rio de Janeiro, RJ, Brazil.

5 – Laboratório de Vírus Respiratório e do Sarampo, IOC, Fiocruz, Rio de Janeiro, RJ, Brazil.

7 – Instituto Nacional de Infectologia Evandro Chagas, Fiocruz, Rio de Janeiro, RJ, Brazil

8 – Instituto D'or de Pesquisa e Ensino, Rio de Janeiro, RJ, Brazil

6 - National Institute for Science and Technology on Innovation in Diseases of Neglected Populations (INCT/IDNP), Center for Technological Development in Health (CDTS), Fiocruz, Rio de Janeiro, RJ, Brazil.

***Correspondence footnote:**
Thiago Moreno L. Souza, PhD

************************************

Fundação Oswaldo Cruz

Centro de Desenvolvimento Tecnológico em Saúde (CDTS)

Instituto Oswaldo Cruz (IOC)

Pavilhão Osório de Almeida, sala 16

Av. Brasil 4365, Manguinhos, Rio de Janeiro - RJ, Brasil, CEP 21060340

#### Additional Results *in silico*

*Molecular Dynamics*

The enzyme-LPV complex was stable with a RMSD value not exceeding 3 Å, but when considering LPV alone, its RMSD exceeded 4 Å over the entire simulation (Figure S1A). By contrast, when complexed with ATV, the RMSD of SCV2-MP reached as much as 4 Å in the last 20 ns of the simulation, while the RMSD of ATV alone did not exceed 3 Å over the 100 ns (Figure S1B). After the step of energy minimization, at the beginning of the molecular dynamics, LPV presented 4 hydrogen bonds with the amino acids Thr25, Ser46, Gly143, and Gln189 (Figures S2A and S3A). After 100 ns of molecular dynamics, LPV interacted with Ser46 HIS41 (Figures 3A and 4A).

ATV's interaction, after energy minimization, occurred with residue Gln189 (Figure S2B and S3B), but after 100 ns of molecular dynamics, Glu166 acted as a hydrogen acceptor and His164 as a hydrogen donor (Figure 3B and 4B).

### Tables

**Table S1**. Interactions from Atanazavir (ATV) with the residues from the active site.

| Docking Interactions of ATA | | |
|---|---|---|
| Ligand | Residues | Distance, Å |
| ATV:H32 | ASN142:OD1 | 1.91 |
| ATV:H32 | HIS164:O | 2.64 |
| ATV:H8 | GLU166:OE1 | 2.44 |

**Table S2**. Interactions from Lopinavir (LPV) with the residues from the active site before (*) and after MD.

| Molecular Dynamics Interactions of LPV | | |
|---|---|---|
| Ligand | Residues | Distance, Å |
| LPV:O1 | THR25:HG1 | 1.99* |
| LPV:O2 | SER45:HG | 1.75* |
| LPV:O | GLY143:H | 2.31* |
| LPV:O2 | GLN189:HE21 | 1.95* |
| LPV:O2 | SER46:HD | 1.70 |
| LPV:O1 | HIS41:HE2 | 2.03 |

**Table S3**. Interactions from Atanazavir (ATV) with residues from the active site before (*) and after MD.

| Molecular Dynamics Interactions of ATV | | |
|---|---|---|
| Ligand | Residues | Distance, Å |
| ATV:H28 | GLN189:OE1 | 2.04* |
| ATV:H31 | HIE167:O | 1.87 |
| ATV:H2 | GLU166:O | 2.51 |

**Legend for the Figures**

**Figure S1**. Root Mean Square Deviation (RMSD) along 100 ns of MD simulation of the Mpro (black) when complexed with A) LPV (red) and B) ATV(red). (A) LPV (red) is rather unstable while Mpro (black) remains stable. In (B), it is Mpro (black) that is destabilized while ATV (red) remains stable along the MD simulation.

**Figure S2**. Representative images of the Root Mean Square Deviation (RMSD) along 100 ns of molecular dynamics (MD) simulation. LPV (A) and ATV (B) are positioned in the Mpro active site at the beginning of 100 ns simulation.

**Figure S3**. Two-dimensional (2D) representation of the interactions of LPV (A) and ATV (B) into Mpro's active site at the beginning of 100 ns RMSD.

Figure S1

A

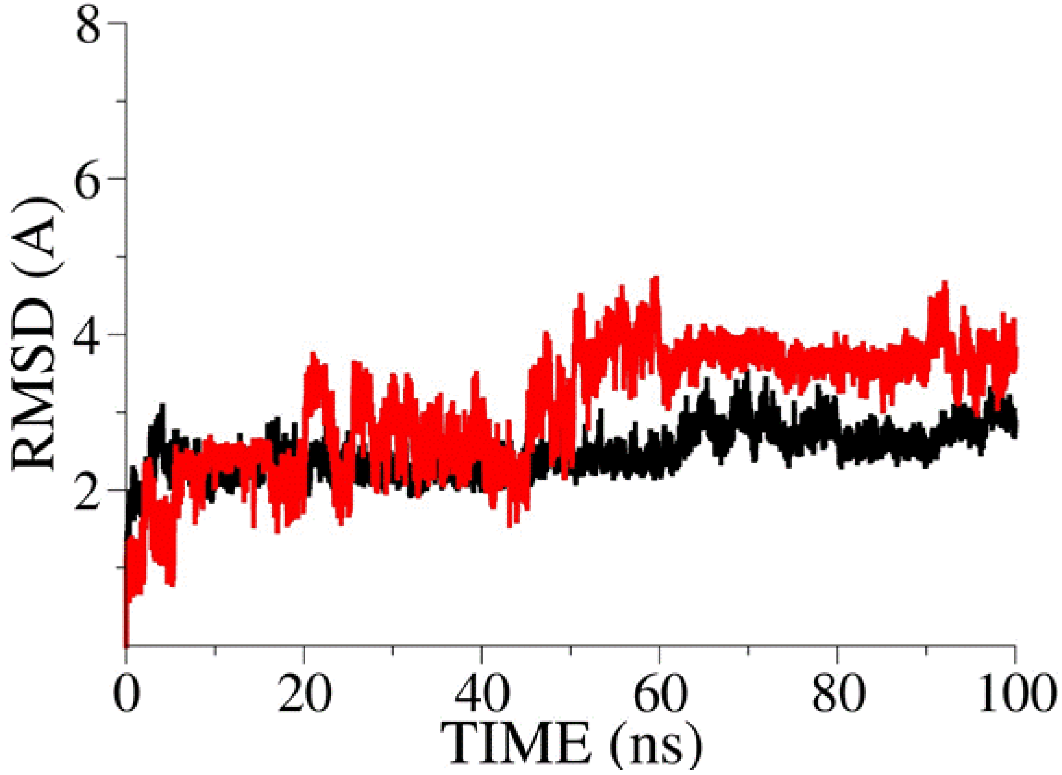

8
6
4
2
RMSD (A)
0
20
40
60
80
100
TIME (ns)


B

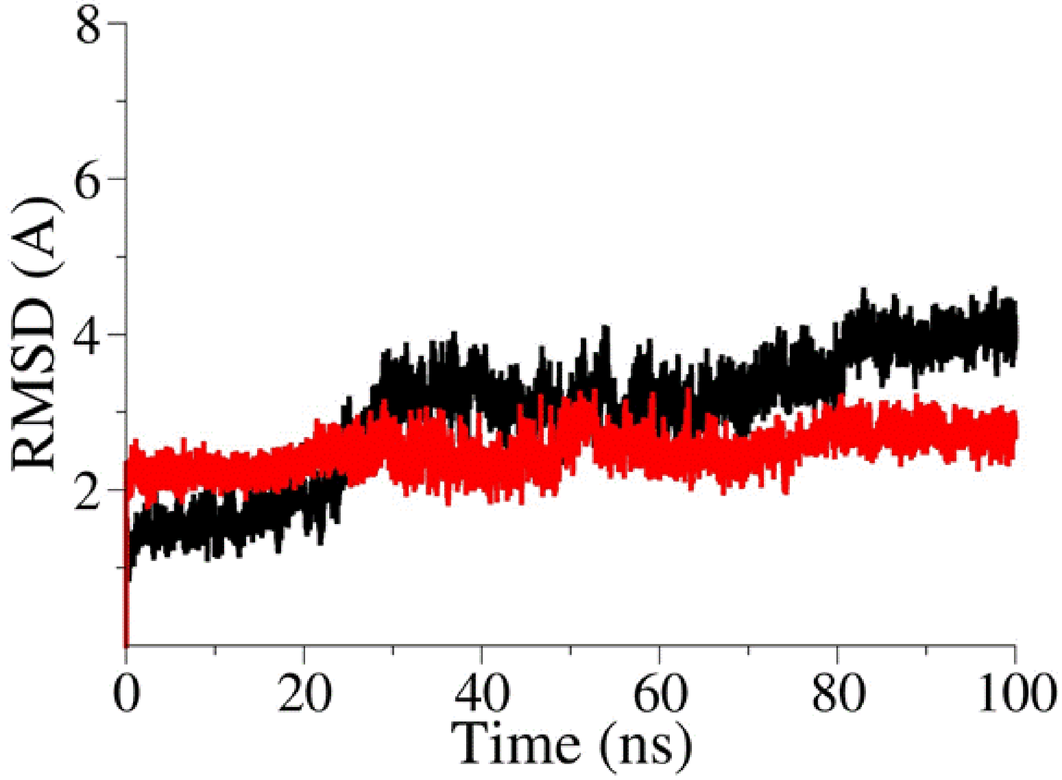

8
6
4
2
RMSD (A)
0
20
40
60
80
100
Time (ns)

Figure S2

A

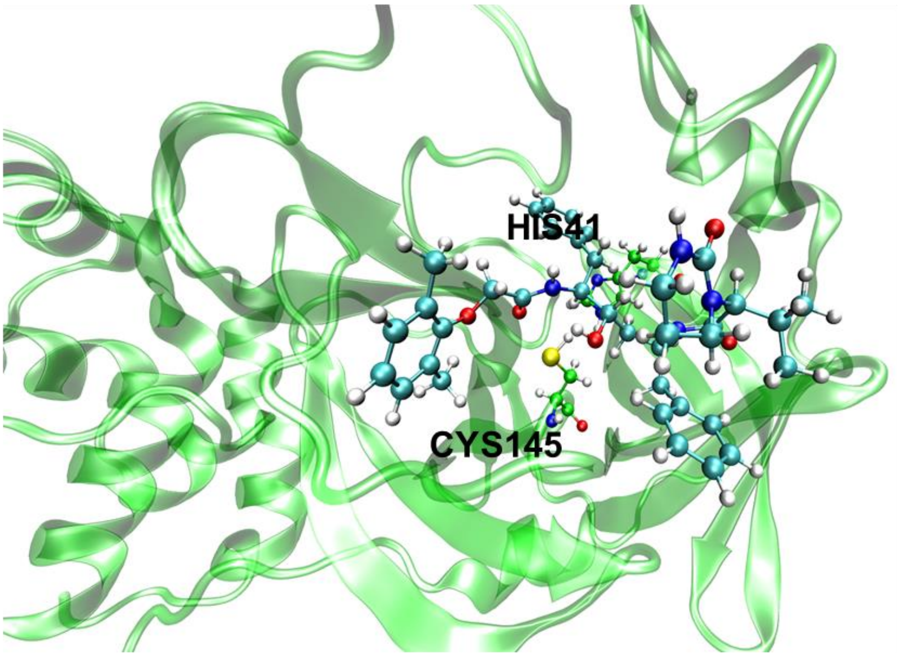


B

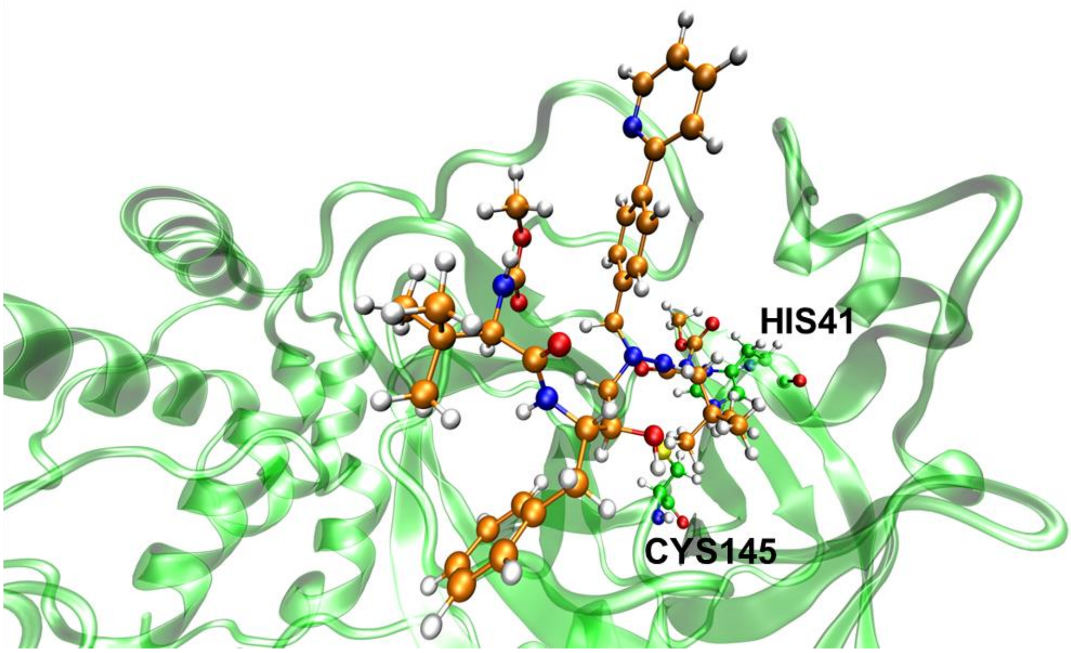

Figure S3

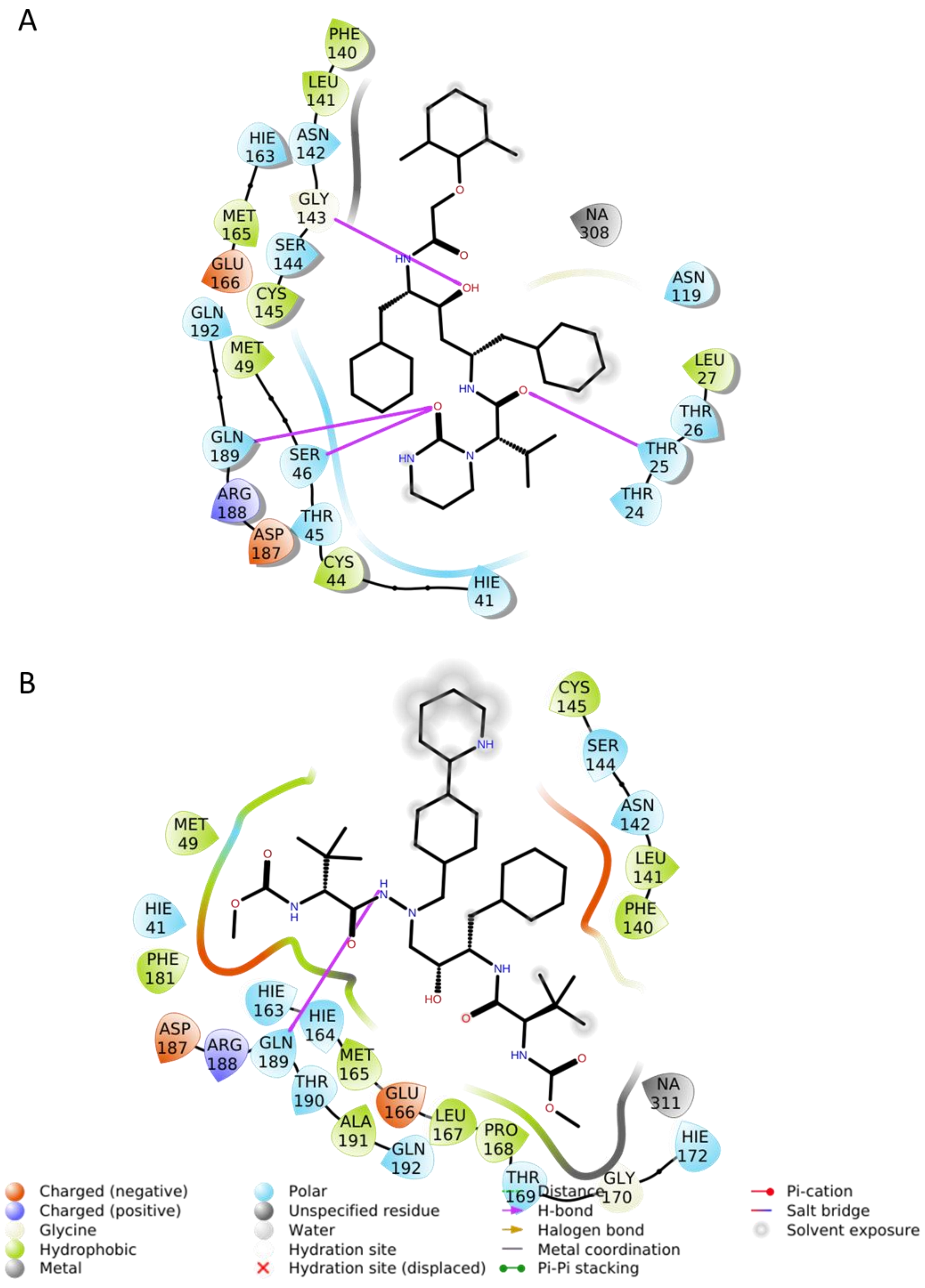
A
PHE 140
LEU 141
ASN 142
HIE 163
GLY 143
MET 165
SER 144
GLU 166
CYS 145
GLN 192
MET 49
GLN 189
SER 46
ARG 188
THR 45
ASP 187
CYS 44
HIE 41
NA 308
ASN 119
LEU 27
THR 26
THR 25
THR 24
B
CYS 145
SER 144
ASN 142
LEU 141
PHE 140
MET 49
HIE 41
PHE 181
HIE 163
HIE 164
ASP 187
ARG 188
GLN 189
MET 165
THR 190
GLU 166
ALA 191
LEU 167
PRO 168
GLN 192
THR 169
GLY 170
NA 311
HIE 172
Charged (negative)
Charged (positive)
Glycine
Hydrophobic
Metal
Polar
Unspecified residue
Water
Hydration site
Hydration site (displaced)
Distance
H-bond
Halogen bond
Metal coordination
Pi-Pi stacking
Pi-cation
Salt bridge
Solvent exposure
